# Finding Antibodies in Cryo-EM densities with CrAI

**DOI:** 10.1101/2023.09.27.559736

**Authors:** Vincent Mallet, Chiara Rapisarda, Hervé Minoux, Maks Ovsjanikov

**Affiliations:** LIX, Ecole Polytechnique, IPP Paris; Sanofi

## Abstract

Therapeutic antibodies have emerged as a prominent class of new drugs due to their high specificity and their ability to bind to several protein targets. Once an initial antibody has been identified, an optimization of this hit compound follows based on the 3D structure, when available. Cryo-EM is currently the most efficient method to obtain such structures, supported by well-established methods that can transform raw data into a potentially noisy 3D map. These maps need to be further interpreted by inferring the number, position and structure of antibodies and other proteins that might be present. Unfortunately, existing automated methods addressing this last step have a limited accuracy and usually require additional inputs, high resolution maps, and exhibit long running times.

We propose the first fully automatic and efficient method dedicated to finding antibodies in cryo-EM densities: `CrAI`. This machine learning approach leverages the conserved structure of antibodies and exploits a dedicated novel database that we built to solve this problem. Running a prediction takes only a few seconds, instead of hours, and requires nothing but the cryo-EM density, seamlessly integrating in automated analysis pipelines. Our method is able to find the location of both Fabs and VHHs, at resolutions up to 10Å and is significantly more reliable than existing methods. It also provides an accurate estimation of the antibodies’ pose, even in challenging examples such as Fab binding to VHHs and vice-versa. We make our method available as a ChimeraX[44] bundle. ^1^

---

Since the first monoclonal antibody entered the clinic in 1986 [19], antibody-based therapeutics have made considerable progress. With over a hundred compounds approved and forty just in the last three years [33, 55], the use of antibodies currently appears as one of the most promising approaches for designing new treatments for patients. Antibody-based therapeutics rely on the identification of antibodies that can bind to a target molecule (antigen) with high specificity through their tips called the Complementarity Determining Regions (CDRs). Monoclonal antibodies (mAb) are the most widely used antibodies. They are composed of two domains called the Antigen Binding Fragment (Fab) and a central stalk that binds to the Fc receptor. More recently, antibody fragments consisting of a single variable antibody domain, engineered from heavy-chain antibodies generated by camelids [1] (nAbs or VHHs) - have attracted interest as an alternative to mAbs [17, 30]. Whether for mAbs or VHHs, initial hits are typically found using immunization [29, 35] or phage display [41, 54]. Before entering clinical studies, initial hits need to be optimized with regards to several properties including their efficacy, manufacturability and safety. Among those properties, the specific binding to the antigen is a critical objective scrutinized from the early phases of the process until drug candidate selection. Binding optimization relies on obtaining the structure of the initial hit since the knowledge of the atomic coordinates at the contact points between the Ab and its target improves the understanding of its mode of action and guides the optimization of the binding affinity [12].

Cryogenic Electron Microscopy (cryo-EM) has become the most common way to experimentally obtain protein structures of therapeutic antibodies bound to their target. In cryo-EM, the target protein is embedded in ice and exposed to an electron beam, resulting in raw noisy images of individual particles. These raw 2D images are then aligned and transformed into a 3D electron probability density [48, 52] from which atomic coordinates are inferred. Recent advances in data collection hardware and software, along with improved data processing pipelines have increased data output. As a result, more academic labs and global pharmaceutical industry have adopted the technology[42]. Lately, a new data collection workflow has been shown to produce 3-4Å structures of a pharmaceutically relevant target protein with 1 hour of instrument time, thus allowing the theoretical resolution of 24 structures a day [15]. However, this raw data needs to be processed from micrographs to molecular structures. While data collection is rapid, current data *processing pipelines* rely on significant manual intervention for simple tasks and decisions, and ultimately take days and weeks to complete.

The recent rise of artificial intelligence-based methods for protein structure prediction holds promise but cannot yet accurately model of the interaction between antibodies and their epitope. Current methods fail to accurately model the diverse CDR loops that are critical for antigen binding [27] calling for significantly more structural data. Unfortunately, the reliance of existing methods on manual intervention significantly hinders cryo-EM density analysis at scale, required for such understanding.

As automation has allowed X-ray crystallography to become a key technique in the structure-based drug discovery pipeline [4], so it should for cryo-EM, making it cheaper and faster, and freeing the time of researchers from button pressing tasks, to structure interpretation and drug engineering. The process of automation is being accelerated by the application of machine learning to the different stages of the pipeline, from data acquisition [5, 21], to preprocessing of micrographs [51], particle picking [2, 58, 59, 60, 61], 2D class selection [34] and 3D heterogeneity deconvolution [40, 62].

Unfortunately, the last step of attribution of the map (fitting of atomic coordinates of a protein into the density), remains a tedious analysis bottleneck. It is still a largely manual process typically done using ChimeraX [43], followed by the local optimization of atomic coordinates using Coot [20]. Some techniques have been developed to help automate this process [37], although with limited accuracy. Furthermore, the methodological challenges associated with this problem have so far impeded the automation of this step using existing machine learning approaches. Specifically, the data is noisy and heterogeneous and the output is high dimensional, making off-the-shelf computer vision methods irrelevant. Tools based on machine learning to trace the sequence in the density were recently developed with good results [28, 46]. They are, however, limited to resolutions better than 4Å and can exhibit prohibitive running times.

In the context of using cryo-EM for optimization of therapeutic Abs, we aim to address the problem of finding Abs (Fabs and VHHs) in cryo-EM maps. To achieve this, we propose `CrAI`, the first fully automatic and efficient machine learning based approach that is applicable at all resolutions better than 10Å without any additional inputs beyond the density. To develop our solution, we introduce a customized deep learning technique, which takes into account and exploits the structural properties of this problem setting. In particular, we leverage the conserved structure of Abs as a prior information[14] to formulate our problem as a special instance of 3D object detection [49, 64]. We gather a novel database of aligned Ab structures and densities, and use it to train a model with a custom loss that involves optimal transport supervision. We test our tool on a set of 215 maps of various resolutions, containing 374 Fabs and 86 VHHs. We successfully find Abs in over 90% of systems, outperforming existing methods by a margin of 25%, while exhibiting **one thousand times** speedup. We make our tool available as a ChimeraX bundle to facilitate adoption.

## Results

### Overview

`CrAI` detects antibodies in cryo-EM maps using a customized deep learning-based technique, trained on our curated dataset comprised of 1430 cryo-EM maps containing Fabs and VHHs. When designing the model, we introduce a custom representation of the structure of antibodies to facilitate the learning process. Specifically, we leverage the *conserved nature* of antibodies to approximate the detailed structure of the output by its *position* and *orientation*. Moreover, we parameterize the orientation in a biologically relevant way that favors the prediction of the CDRs location. Finally, the list of such representations of antibodies for a system is transformed into a grid overlaid over the cryo-EM map, such that the position of an antibody is encoded as an offset from a grid cell. The encoding of the output is shown in Figure 1A.

**Figure 1:**
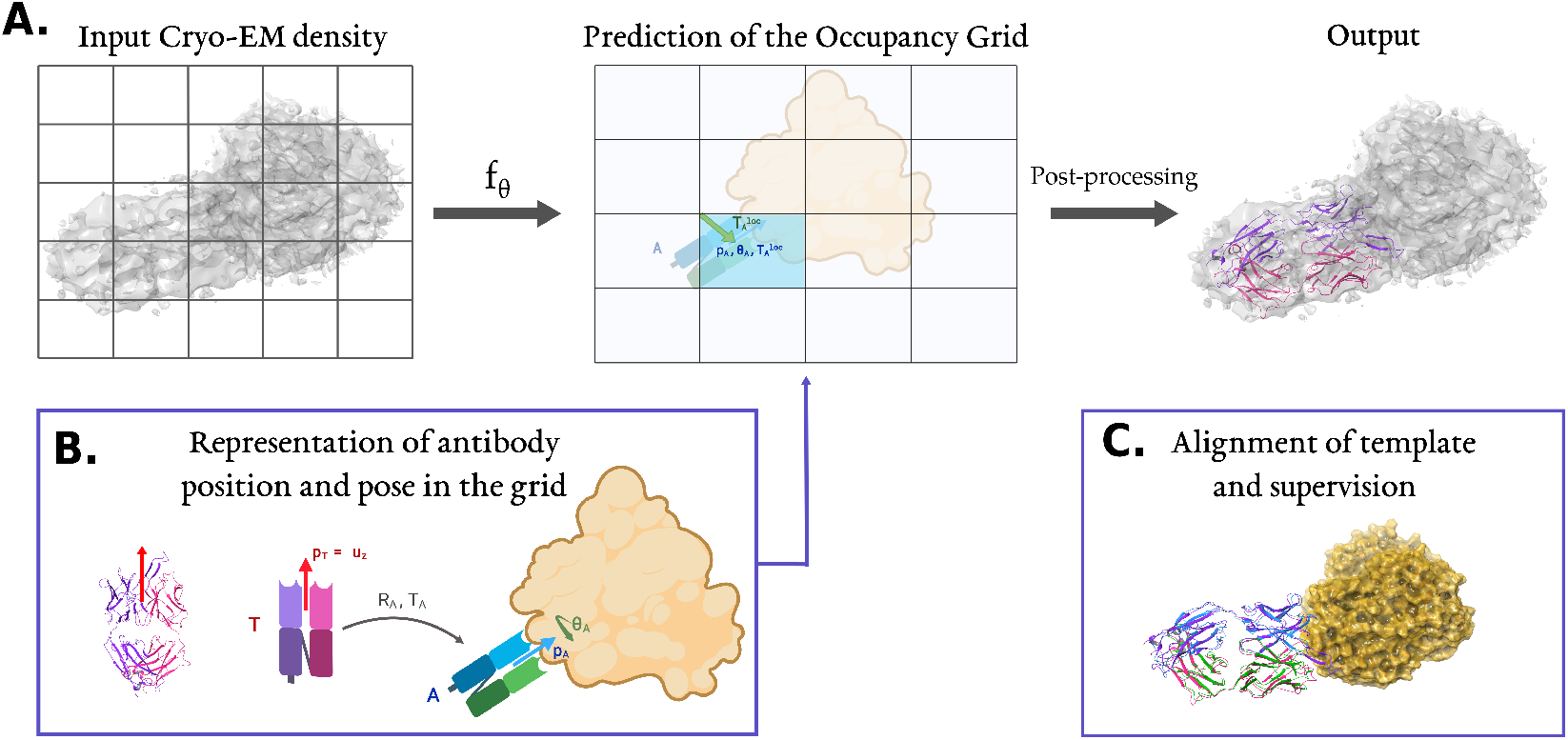
**A**. `CrAI` predicts an occupancy grid that represents antibodies found in an input density. The prediction of this occupancy grid can be post-processed into a PDB containing the predicted antibodies’ structures. **B**. The atomic structure of our template (pdb `7lo8`) is displayed next to its cartoon, with 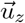 shown in red. We compute optimal alignments 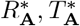 of our template onto Abs and decompose the rotation into 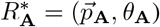.We decompose the translations into a position in a grid 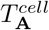 and an offset from the grid corner 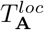. We thus obtain a grid with zero values except for cells containing an Ab, in those cells, we have 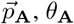 and 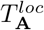. **C**. Example alignment of our template (*red, purple*) with the experimental structure of system `6bf9`: antigen (*orange*) and Fab (*blue, green*). As can be seen, our template aligns well to other antibodies (RMSD = 1.8Å).

We use an architecture that consists of a 3D UNet trained to predict this output representation from the input map using a custom loss with two components. The first one enforces the prediction of the location of grid cells containing an antibody, and relies on optimal transport to take into account the geometric aspect of the prediction. The other part controls the prediction of the pose and the discrimination between Fabs and VHHs. Once the model makes a prediction, persistence diagrams [6, 7] are used to select the relevant results which are transformed into a PDB file. Remarkably, this procedure does not require anything beyond the input density. Our pipeline is illustrated in Figure 1.

### Baselines and Metrics

To the best of our knowledge, no tool enables predicting the position of antibodies solely from a density map. We benchmark against `dock_in_map` [37], a tool that takes the density along with the known atomic protein structures that need to be docked in the map. Considering that at test time, one does not have access to the ground truth structure, we ran `dock_in_map` with a fixed template Fab or Fv. Additionally, we ran `dock_in_map` with the actual structure giving us an upper bound of its performance.

Independent models were trained on the training sets obtained with the `random` and `sorted` splits (see Methods Section). We make inference of those models on their respective test sets, providing the network with the number of Abs to find (denoted as `num`) or relying on automatic thresholding to infer this number (`thresh`). For each system, a prediction method results in one or several predicted Ab positions. When evaluating different approaches, individual predicted antibody positions need to be matched with an actual antibody position. This matching process is accomplished using the Hungarian algorithm [36] on the distance between the center of mass of predicted and actual antibodies.

The distribution of distances between centers of mass of the predictions and experimental structures shows that we outperform `dock_in_map`, even in the idealized scenario that uses the ground truth structure (see Supplementary Figure S.2). In the following, we will compare to this idealized and more challenging scenario as our baseline. Moreover, we observe that the predicted distances are bimodal: a first peak corresponds to systems predicted successfully and another spread mode corresponds to failed prediction. Hence, we will report our results in terms of **hit rates**: a prediction is deemed as a hit if it is closer than 10Å to the ground truth. As can be seen in the histogram, performance is not very sensitive to the choice of this threshold. We report our results for individual Abs (`ab`) as well as aggregated by systems (`sys`) so that a system with many Fabs does not influence the results significantly.

### CrAI accurately finds Abs

Our key results of Fab detection are presented in Table 1. In the first two rows, note that `CrAI` drastically outperforms `dock_in_map` in terms of hit rates. This holds true in all settings for an overall hit rate going from approximately 69% to 95%. In terms of distance between the centers of mass prediction and the ground truth, both methods give results below 2.5Å which is close to optimal considering the map resolutions. Given a successful hit, `dock_in_map` appears to be a bit more precise, which is expected as it uses the ground truth structure instead of a template. Moreover, distances are computed only over successful systems and thus include *25% more systems in the* `CrAI` *column*. One could first rapidly screen a map with a high hit rate using `CrAI`, then precisely refine results in a density neighborhood using ChimeraX fast local refinement tool, *fitmap*. This possibility is enabled in our ChimeraX bundle.

**Table 1:** Fab detection performance of the performance of the benchmark tool `dock_in_map` and of `CrAI`, provided with the ground truth number of objects (`num`) or not (`thresh`). We report the Hit Rate (defined as the number of prediction below a threshold of 10Å) on different data splits (`random` or `sorted`), aggregated by systems (`sys`) or not (`ab`). We additionally report the mean distance of these successful predictions, and the mean runtime of the tool.

| Method | Hit rate |  |  |  |  |  | Hit distance | Runtime |  |
| --- | --- | --- | --- | --- | --- | --- | --- | --- | --- |
|  | random |  | sorted |  | mean |  |  | CPU | GPU |
|  | sys | ab | sys | ab | sys | ab |  |  |  |
| dock_in_map | 66.6 | 61.9 | 71.3 | 66.0 | 69.0 | 64.0 | **0.73A** | 883s | - |
| CrAI | **97.3** | **96.7** | **96.9** | **95.7** | **97.1** | **96.2** | 1.73A | **1.9s** | **0.47s** |
| CrAI thresh | 95.9 | 95.2 | 92.7 | 92.0 | 94.3 | 93.6 | 1.71A | **1.9s** | **0.47s** |

### CrAI can be used with unknown number of Fabs

We now evaluate `CrAI` in the scenario where it also automatically estimates the *number* of Abs, and thus the network is given *only the density* without any additional information. As shown in the second and third lines of Table 1, we retain most of the performance even in this more challenging, and more realistic setting. We find this remarkable, since the densities are highly heterogeneous and contain between one and six Abs. This is to compare to the result of `dock_in_map` that is additionally given the *experimental structures of all antibodies* that are to be found in the map. In contrast, the fact that our approach can adapt to different scenarios strongly highlights its effectiveness and flexibility, and sets it apart from *any* existing method in terms of its ability to detect antibodies in Cryo-EM densities in a fully automatic manner. Furthermore, as highlighted below, our approach is also orders of magnitude faster than the baselines, making it the first truly scalable and practical method for efficiently detecting Fabs in Cryo-EM densities.

### CrAI runs fast, at all resolutions

The average runtime of `dock_in_map` on our validation set is a prohibitive 883s/system. This mean is optimistic because we stopped a few systems after 5 hours of computations (letting them run longer would increase the mean). In comparison, our tool runs in **0.47s/system**, i.e., more than a thousand times faster when using a GPU (A40). Even when using only one CPU, our tool runs in 1.9s/system, four hundred times faster than `dock_in_map` and fast enough for this computational step to integrate seamlessly in an analysis pipeline. This stems from the complexity of our algorithm, that is linear with regards to the grid size.

Supplementary Figure B.2 shows how our performance depends on resolution of the input map, over all of our data sets. Contrary to `dock_in_map`, we do not see a correlation and thus claim that `CrAI` is *robust to low resolution*. This is a novel result as machine learning methods for tracing such as ModelAngelo [28] only work for resolutions better than 4Å.

### CrAI finds VHHs

We now turn to the prediction of VHHs. They are more challenging for a machine learning model, because we have less training data (278 vs 722 training examples). Moreover, the binding modes of VHHs are less canonical which can affect the decomposition of our rotation. Finally, VHHs are smaller and tend to be more buried into the density - as opposed to Fabs often sticking out. Our results are presented in Table 2.

**Table 2:** Performance at detecting VHHs. The convention used is the same as above.

| Method | Hit rate |  |  |  |  |  | Hit distance |
| --- | --- | --- | --- | --- | --- | --- | --- |
|  | random |  | sorted |  | mean |  |  |
|  | sys | ab | sys | ab | sys | ab |  |
| dock_in_map | 77.8 | 68.8 | 90.5 | 89.1 | 84.2 | 79.0 | **0.5Å** |
| CrAI num | **100** | **100** | **90.9** | **90.6** | **95.5** | **95.3** | 1.66Å |
| CrAI thresh | 99.1 | 98.7 | 84.1 | 81.2 | 91.6 | 90.0 | 1.57Å |

We achieve a positive result with a 15% hit rate boost over `dock_in_map` overall, and 9% in the `thresh` setting. Beyond the intrinsic difficulties outlined above, we underline two additional factors to explain why the performance gap is smaller than for Fabs. On average, `dock_in_map` performs much better for VHHs than for Fabs, which could stem from their better resolutions. Moreover, on the `sorted` split, it outperforms itself (performance over the whole data is similar to *random*), which can be explained by the variance of estimators on small sized test sets. Nevertheless, we emphasize that `CrAI` still outperforms `dock_in_map` in terms of the number of successful detections, without relying on any additional side (ground truth) input and being orders of magnitude faster.

### Ablation study

To assess the relevance of different design choices of our approach, we retrain several models without specific individual features in our design. Since this means training a new model every time, we only perform this analysis in the Fab `random` split setting. We report the performance in the `sys` setting, but results are consistent in the `ab` setting. We try replacing Persistence Diagrams (PD) with a naive Non-Maximal Suppression (NMS) [22, 25] that amounts to zeroing predictions around local minima. We also trained our model without using the Optimal Transport (OT) component, as in done in most classical object detection approaches [24, 50]. We also tried to keep the method fixed, but to disrupt the template encoding by decomposing the rotation using *u*_*y*_ as the main vector. We present the results in Table 3.

**Table 3:**
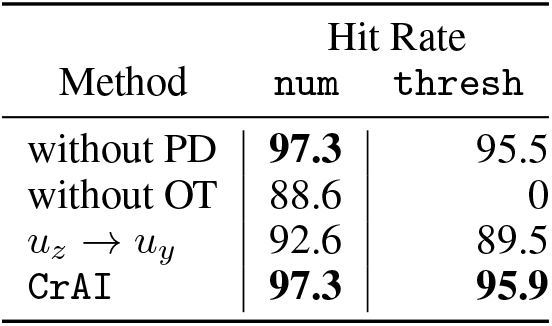
Hit rates for different ablations of our method, on the Fab random split, in the sys setting.

| Method | Hit Rate |  |
| --- | --- | --- |
|  | num | thresh |
| without PD | **97.3** | 95.5 |
| without OT | 88.6 | 0 |
| $u_z \rightarrow u_y$ | 92.6 | 89.5 |
| CrAI | **97.3** | **95.9** |

Persistence diagrams seem to enhance results in the `thresh` setting, with a limited impact. However, when removing optimal transport, performance collapses especially on the ability to predict the number of objects. This can be explained considering that antibodies cannot overlap (in contrast to detecting pedestrians in images for instance). Optimal transport helps to attribute a single detection to a region rather than enabling an arbitrary number of potentially overlapping/conflicting detections. Training the model with *u*_*y*_ instead of *u*_*z*_ also significantly weakens detection performance.

### CrAI finds meaningful poses

After validating the position of our predictions, we consider the predicted poses: are Abs in the correct orientation? Using the decomposition of rotations into predicting a vector 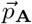 and an angle *θ*_A_, we can compute the angles between predicted values and actual ones. We provide histograms for the distribution of these angles in Figure 2, to show that most systems are predicted accurately. For the Fab data, the angle between 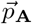 and its prediction is on average of 7.8° and the one for *θ*_A_ is 11.0°, low enough values to make the prediction almost overlap with ground truth. For the VHH data, those values are on average 7.5° and 9.7° respectively. Finally, the model trained using 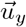 instead of 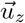 has an angle error of respectively 10.2° and 11.6°, significantly higher on the vector prediction. Hence, predicting 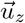 is easier than 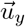, which validates the non-restrictive inductive bias that we introduced in our formulation.

**Figure 2:**
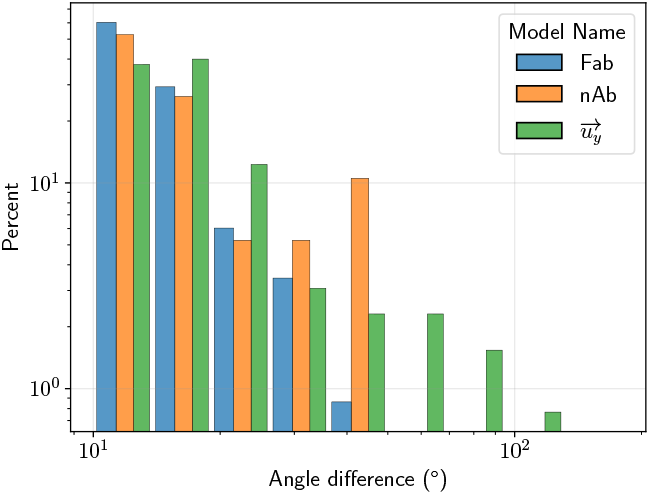
Histograms of the angles between the predicted and experimental 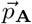 vector for Fabs, VHHs and 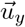 model

### Further analysis

#### Case studies

Detailed evaluation results for 6 systems of the random split test set are shown in Figure 3. Those six systems were picked because of their relevance in the context of drug discovery, at resolutions ranging from 3 to 8Å, include four Fabs whose location was found by our tool consistently and correctly. `dock_in_map` on the other hand, only finds the right location in one case, predicts a shifted location on another and completely fails its prediction on the last ones. The two remaining systems, with resolutions of 3.1 and 3.8Å respectively, contain VHHs for which `CrAI` detects the location correctly and for which `dock_in_map` fails. There were no examples in which dock in map succeeded and `CrAI` failed. Of the 6 systems shown, 4 used Chimera for manual docking, one doesn’t mention the docking program and one uses MDFF[56], a molecular dynamics software not commonly employed by experimental structural biologists. It is hard to evaluate ChimeraX placement in the map as the time and the success of placement isn’t automatic and it is user dependent. For the lower resolution structures, chain building programs are not suitable as resolutions better than 3.5 Å are needed.

**Figure 3:**
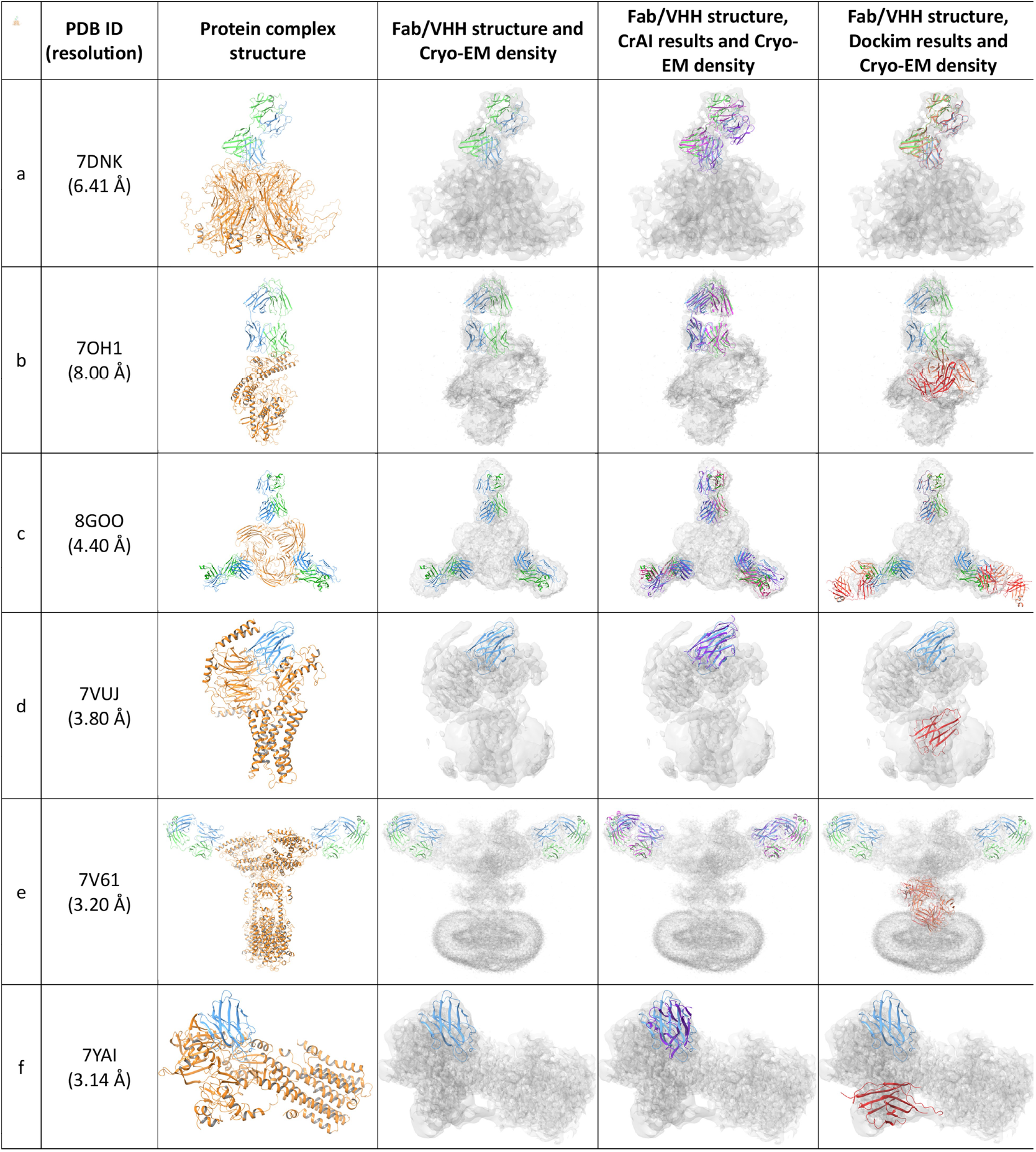
In this Table, columns correspond to 1) PDB code and EM resolution; 2) Protein structures from the PDB (the antigen is displayed in orange, the heavy chain in blue and the light chain in green); 3) EM map and its corresponding structure; 4) Superimposition of the Fab or the VHH from the PDB structure (blue/green) with the `CrAI` results (purple); 5) Superimposition of the FAB or the VHH from the PDB structure with the `dock_in_map` results (red). The rows correspond to six systems of our test set: (a) Human papillomavirus type 58 pseudovirus (HPV58) in complex with the Fab fragment of 5G9 at a resolution of 6.41Å, (b) Tetanus neurotoxin LC-HN domain in complex with TT110-Fab1 at a resolution of 8.00Å, (c) Structure of beta-arrestin2 in complex with a phosphopeptide corresponding to the human C5a anaphylatoxin chemotactic receptor 1 C5aR1 at a resolution of 4.40Å, (d) Cryo-EM structure of a protein complex containing a class A orphan GPCR (GPR139), stabilized by a VHH (Nb35) at a resolution of 3.80Å, (e) ACE2 -Targeting Monoclonal Antibody as Potent and Broad-Spectrum Coronavirus Blocker at a resolution of 3.2Å, (f) Cryo-EM structure of SPCA1a in E1-Ca-AMPPCP state subclass 3 at a resolution 3.14Å.

The first two examples are low resolution illustrations of the structural characterization of neutralizing antibodies, in the context of cervical cancer for the Human papillomavirus type 58 pseudovirus (HPV58) in complex with the Fab fragment of 5G9 at a resolution of 6.41Å [26](Figure 3a) and the viral infection with tetanus for the neurotoxin LC-HN domain in complex with TT110-Fab1 at a resolution of 8.00Å [47] (Figure 3b). While the low resolution of the first example does not preclude both `CrAI` and `dock_in_map` to correctly predict the position of the Fab, at 8.0 Å, only `CrAI` correctly positions the Fab. Membrane proteins are notoriously difficult to characterize structurally and are often solved at lower resolutions than soluble ones. Structural information is important to decipher the signaling and regulatory mechanisms of this class of proteins. G protein-coupled receptors (GPCRs) in Figure 3c and d play a crucial role in signaling and are one of the major target family of drugs. Three Fabs are present in the density of beta-arrestin2 (Figure 3c) in complex with a phosphopeptide corresponding to the human C5a anaphylatoxin chemotactic receptor 1 C5aR1 at a resolution of 4.40Å [39]. `CrAI` correctly finds all of them while `dock_in_map` misses to correctly predict two out of three. In Figure 3d, the class A orphan GPCR (GPR139), [63] is captured with and stabilized by Nb35 in different conformational states highlighting allosteric modulation roles of VHHs in GPCRs. In this first example with a VHH in a protein complex, `CrAI` correctly finds the location of the VHH while `dock_in_map` failed. In the fight against covid-19, a search for potent and broad-spectrum coronavirus blockers was initiated and many antibodies were shown to block the entry of several variants of SARS-CoV-2 (SARS-CoV-2, SARS-CoV-2-D614G, B.1.1.7, B.1.351, B.1.617.1, P.1, SARS-CoV, and HCoV-NL63) by binding to human ACE2, without causing severe side effects. In Figure 3e, ACE2 is found in complex with targeting monoclonal antibodies at a resolution of 3.2Å [10]. This system illustrates `CrAI` correctly finds the two Fabs in the density, while `dock_in_map` is unsuccessful. The final example in Figure 3f is the Cryo-EM structure of SPCA1a in E1-Ca-AMPPCP state subclass 3 at a resolution 3.14Å [11]. In this case, with a medium resolution, the location of the VHH Nb14 is properly found by `CrAI`, however, its orientation is not correct. When analyzing this protein complex, it appears that the binding mode of Nb14 to its antigen, the Calcium transporting ATPase type 2C member 1, is non-conventional, with the framework region of Nb14 lying on the surface of the antigen while the CDRs are not oriented toward the epitope. The prediction made by `CrAI` followed the conventional binding mode of a VHH, with the CDRs facing the epitope of the antigen. This specific example highlights a potential bias of `CrAI` in correctly predicting the orientation and the binding mode of antibodies and VHH. It can be noted that in this specific case, `dock_in_map` fails to identify the correct location of Nb14.

#### Failure analysis

Since the number of failed systems with `CrAI` is relatively low, we visualized all *all of them* (for the `sorted` split) in Supplementary Figure S.3. After inspecting these failures, we identified two dominant sources of errors as reported in Table 4A. A visual depiction of these two error categories is shown in Figure 4B.

**Figure 4:**
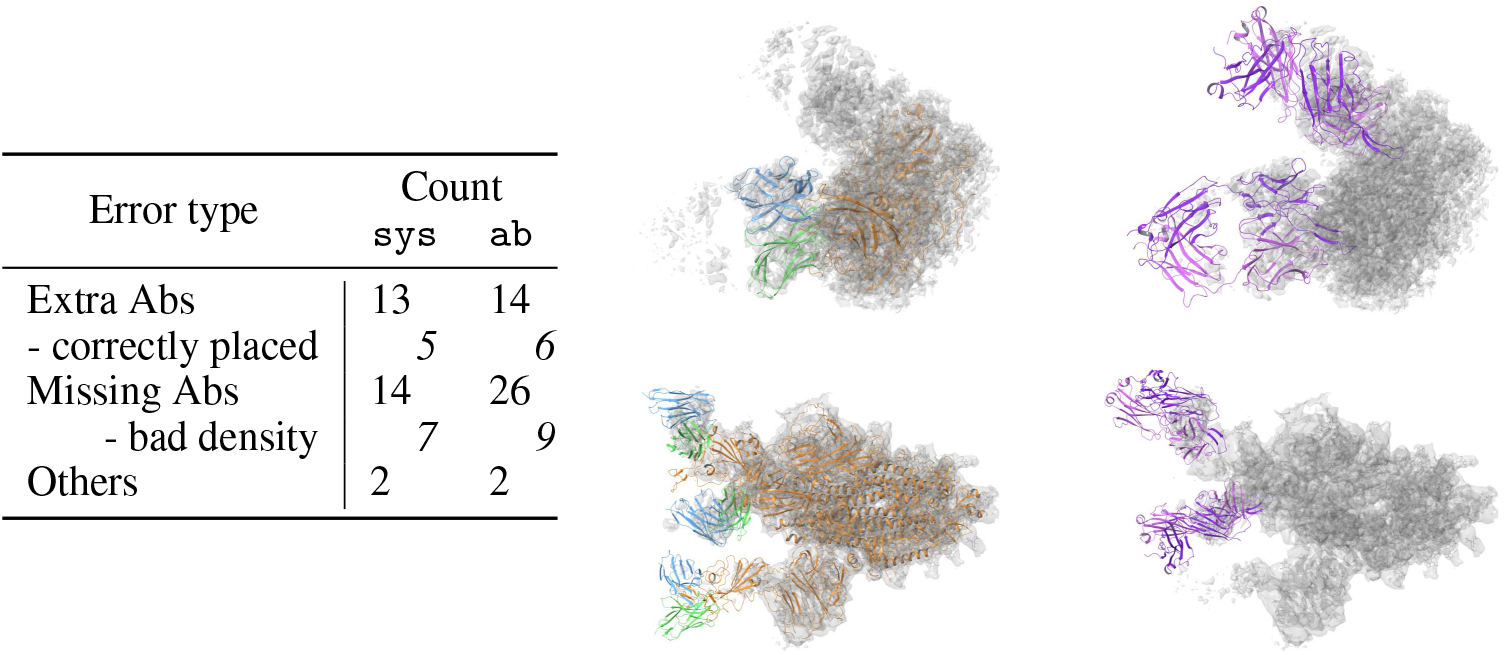
*Left:* An exhaustive classification of our errors on the `sorted` split for both Fabs and VHHs. *Right:* Illustration of the two main modes of failures on two examples. The first row corresponds to `8E8R` and the second to `8HEC`.---Most errors result from predicting more abs - including scenarios where these extra predictions are reasonable like for `8E8R` - or fewer abs - including scenarios where the missing ab falls in a region with very poor density like in `8HEC`.

The vast majority of failures amount to either predicting too many antibodies or not predicting enough.

When predicting too many Abs, interestingly, the right ones are *always* included in our prediction. Moreover, in 5 out of these 15 errors, the extra predictions were found to correspond to an antibody that *actually appears in the density*, such as an element from another asymmetrical unit, and thus can be considered as accurate detections. In the eight remaining cases, two extra predictions result from a wrong ordering (the wrong prediction is ranked first) and four result from a faulty thresholding: the artifacts have lower probabilities but fail to be automatically discarded. Overall, we remark that these additional predictions are both infrequent and can easily be discarded by practitioners.

Furthermore, predictions for 14 systems lack one or more Ab. In over a third of the cases, this could be explained by a poor density of the map around the missing antibody. Forcing `CrAI` to produce additional predictions on those 12 systems, we observe that 16 out of 20 missing Abs can be found, often obtained after just one or two extra predictions. Thus, our software offers the option to predict a certain *number* of predictions instead of relying on automatic thresholding, to allow practitioners to capture potential missing annotations.

In summary, we noticed that most of our errors originate from the ordering and thresholding of the predictions and not from the detection of antibodies.

To further validate this observation, we performed a study of the accuracy of `CrAI` and `dock_in_map` across a varying number of predictions. For this, we forced both methods to output up to ten predictions and then computed the hit rates obtained when keeping the k-best predictions. While our method can easily output ten predictions, to get this result for `dock_in_map` we independently ran it on a growing set of inputs. The first few inputs correspond to the observed structures of antibodies while additional ones are copies of these structures in a random order.

In Figure 5 we plot the fraction of all antibodies captured by different approaches, when making a growing number of predictions per system. The blue curve represents the best achievable results. It does not always equal one as some systems contain multiple antibodies. As can be seen in this figure, a small discrepancy exists between the ground truth and our tool. However this discrepancy disappears around *k* = 6 predictions, suggesting that with this number of detections per system our method captures *all* antibodies. This further justifies that most of our errors originate from the thresholding. Thus, if our tool fails to detect an antibody, practitioners can ask for more predictions with a high chance of seeing it predicted. `dock_in_map` has a much wider gap that tends to stay consistent despite allowing it to output more predictions.

**Figure 5:**
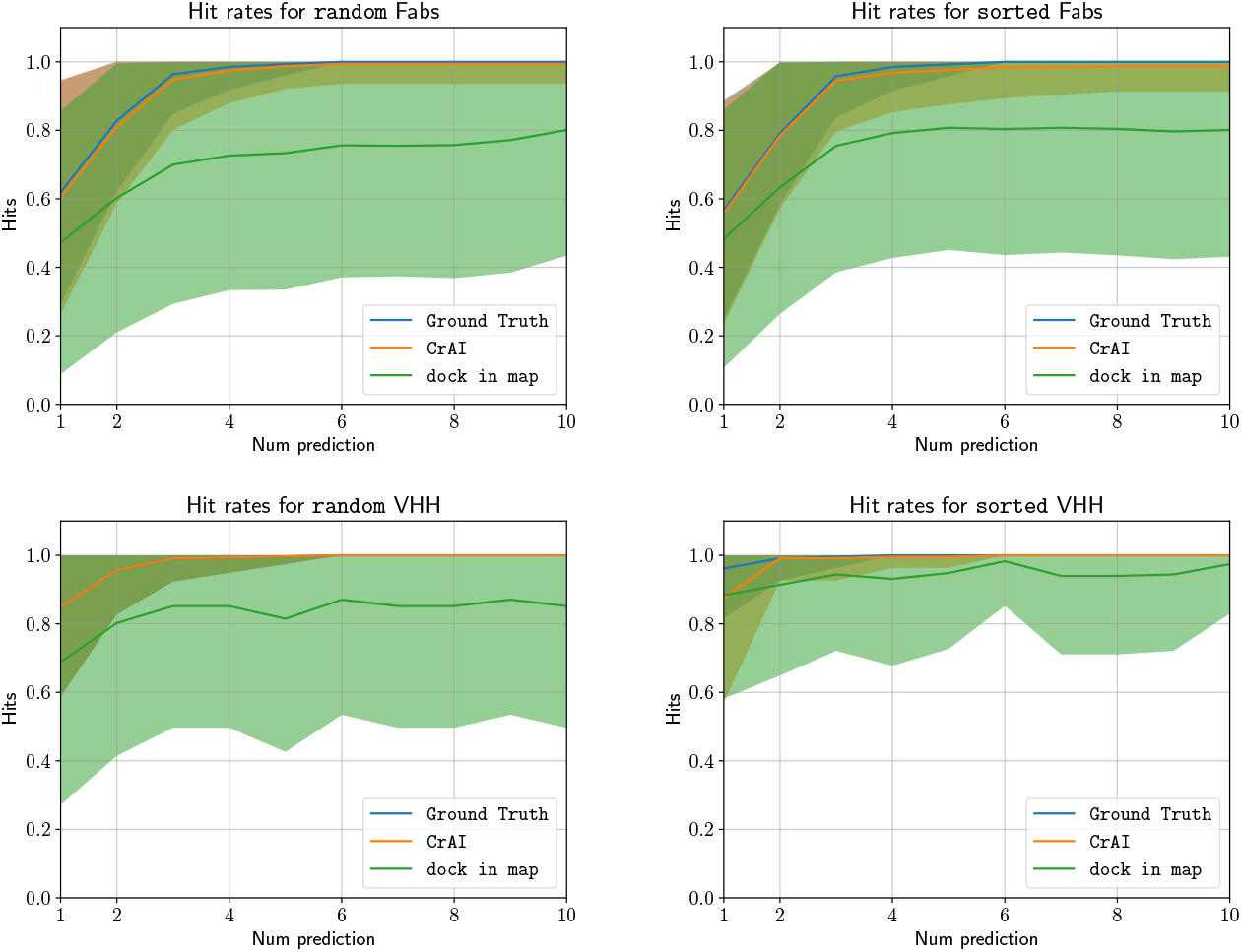
Hits of different approaches (y axis) as a function of the number of predictions (x axis) in our four data settings. The solid lines represent the mean performance of systems and shaded regions correspond to the variance. Even if we observe a small discrepancy between our prediction and the ground truth line, note that most errors originate from the automatic thresholding feature, and our method achieves near perfect hit rate with as few as 6 detections, compared to `dock_in_map`, which fails to capture a significant number of present structures.

## Discussion

In this paper, we addressed the problem of automatically finding the Abs in cryo-EM densities. This step currently constitutes a tedious, manual step thus hindering efficient and scalable structure estimation.

To achieve this goal, we proposed a customized solution, which exploits the structural properties and addresses the specific challenges of the problem, such as handling significant data scarcity and heterogeneity. Specifically, we leveraged the conserved structure of Fabs to cast this problem as an object detection one. We gathered and curated a database to enable a data-driven solution. Finally, we then designed a customized pose representation and loss based on optimal transport, which all help integrate prior information, while remaining efficient and flexible. Using our approach, no extra input is required to predict the number, position and pose of Fabs and VHHs in a density.

We validate our results on experimental maps and find the Ab positions with a hit rate above 90%, which represents a 25% (resp 15%) improvement on Fabs (resp VHHs) over existing methods, while requiring no extra inputs and being *thousands of times faster* even when run on a single CPU. We show that the predicted pose correlates well with experiments, and illustrate our tool’s performance on six systems relevant to drug design.

We believe that this tool can reduce the burden on structural biologists working with cryo-EM densities of Abs and accelerate the resolution of 3D complexes of Abs bound to their antigen. In line with this objective, the method ships as a ChimeraX[43] bundle to enable seamless integration.

In the future, it will be interesting to expand our approach to other conserved families, beyond Fabs and VHHs, and enable placing several folded domains into a density, which has so far been done without machine learning, [8, 37]. More broadly, we could replace a fixed template by a parametrized family of embeddings of the folded domains, expanding the expressive power of our framework.

The exceptional throughput of our tool opens the door to finding Abs in the output of heterogeneous reconstruction or even continuous distribution of maps, and thus to capture several modes of antibody binding. Moreover, the automatization of cryo-EM structure resolution by our technique enables the generation of antibody-antigen complexes at a larger scale, a critical step to improve our understanding and detailed modeling of antibody binding. More structural information will open possibilities for better and more accurate antibody modeling, that will speed up drug discovery process of bio-therapeutics.

## Methods

### Building a database

We build a curated antibody database to train and test our method, comprising Fabs and VHHs. Fabs are composed of one constant and one variable domain for each of the heavy and the light chains. The two variable domains are denoted as the Fv variable fragment (Fv). VHHs are antibody fragments that correspond to the sole variable domain of the heavy chain. They represent a promising family for therapeutic antibodies.

The Fab data is originally fetched from *SabDab*[18] in the form of a list of protein chains. We fix a few broken annotations, notably for systems containing both Fabs and VHHs (more details in Section A.3). Using the PDB[3], we find the corresponding cryo-EM maps and download all corresponding maps and structures. The maps contain densities of both Fabs and Fvs, but the constant region of Fabs is often missing in the deposited structure. To avoid false negatives in our data, we chose to predict the position of the Fv as this is the only part consistently reported. Then, we remove systems with resolution below 10Å or ones with no antibodies or antigen chains, and group all hits pertaining to the same structure together, yielding a total of 1032 systems.

The resulting map files can be enormous (up to 10^9^ grid cells) especially for symmetric assemblies, such as viruses. Since the asymmetric unit only occupies a fraction of the map, using whole maps would create false negatives. Indeed, some regions of the map correspond to proteins omitted in the deposited structure (e.g. pdb `7kcr`). Hence, we crop the original maps around the structure with a margin of 25Å. Additionally, we resample maps to a fixed voxel size of two Å and normalize them by zeroing out negative values and dividing by the maximum of the map.

We split this dataset following a temporal splitting strategy with more recent systems in the test split (`sorted`). While this procedure is often used to provide a realistic use case, it can also introduce a bias, for instance towards COVID structures. Hence, we also report our performance following a random split (`random`). We obtain 722, 155 and 155 systems in the train, validation and test splits respectively. Note that a single system can include several Fabs. On average, there are 2.25 Fabs per system in our dataset. The number of Fabs per split is 1,627, 320 and 374 respectively - with similar numbers for the `random` split. We now have a split database of cropped, resampled, normalized cryo-EM density maps containing Fabs.

All of these steps are repeated to create a VHH database. We started from *SAbDab-nano*[53] and applied the same filters on the raw data, resulting in 398 systems. Then, we performed the *random* and *sorted* splitting procedures as above. This amounts to 278, 60, 60 `systems` in each split and 458, 74, 86 when counted as `ab`, for a mean of 1.55 VHHs per system - again with similar numbers for the `random` split.

### Motivation

The design of our approach, `CrAI`, is motivated by several methodological challenges. First, due to the challenges inherent in the limited scale and significant noise present in the training data, we incorporate the conserved structure of antibodies as prior information to make learning more data-efficient and robust. Second, to facilitate antibody pose finding in the presence of limited data, we use a custom parametrization of the rotation that takes into account an *expected* pose, while being flexible to allow arbitrary orientations. Third, we introduce a fully convolutional design to accommodate arbitrary grid sizes that might be present at inference time. We also train the network with rotation augmentation to approximate rotation equivariance and accommodate arbitrary orientations. Finally, we introduce a custom loss when training our model. Our loss incorporates a formulation based on Optimal Transport along with persistence diagrams to better capture the geometric aspects of our problem, such as predicting non-overlapping objects and including *distance*-based penalties between our prediction and the ground truth. Finally, we introduce a Non-Maximum Suppression step based on persistence diagrams, which selects the most likely individual detections given a predicted probability map.

### Problem formulation

Starting from an input cryo-EM map, we want our method to output the 3D coordinates of one or several Abs. Because of the highly conserved structure of Abs, we can simplify our problem by only predicting how to align a fixed antibody template **T** with the Abs, without deformations. We used the Fab structure in the pdb `7lo8` as a template and computed the optimal alignments with `pymol align`[16] during data pre-processing. We provide an example alignment of our template in Figure 1A.

More formally: Let 𝒳 be the cryo-EM density we consider, *n*_𝒳_ the number of Abs contained in this density and 𝒜_𝒳_ = {**A**_*i*_, 0 ≤ *i* < *n*_𝒳_} the set of such Abs. Given an Ab **A** and a registration objective d, let 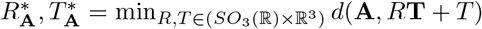 be the translation and rotation that best align **T** to **A**. Finally let 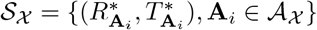 be the set of optimal alignments. Note 𝒮_𝒳_ consists of elements of the Euclidean group in 3D that is six-dimensional, whereas elements of 𝒜_𝒳_ are three-dimensional coordinates for hundreds of atoms. In this paper, we aim to predict 𝒮_𝒳_ instead of 𝒜_𝒳_.

The optimal rotations mentioned above can be parametrized in many ways. Let 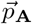 denote the unit vector oriented from the center of mass of an Ab **A** towards its antigen. Since canonical binding tends to happen through the CDRs, we observe that it is easier to predict 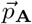 than the rotation around 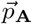 from the density. Therefore, we decomposed 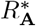 into a rotation transforming 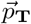 into 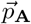 and one 2D rotation around 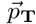 of angle *θ*_𝒜_. The generality of this decomposition is established in Supplemental A.1 and its relevance is shown in Figure 2.

Our problem is now formulated as an object detection problem, which consists of detecting, localizing and aligning the Abs in a given density. Following common practice in the object detection literature [50], we overlay an occupancy grid 𝒢 ^𝒳,S^ of size **S** over our input: this grid contains ones for cells encompassing an Ab and zeros elsewhere. The size of this grid cells corresponds to a fixed spatial location and not on the resolution of the map. We then decompose each translation into two parts: one going to the corner of the occupied cell and one from this corner to the 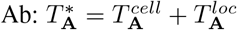, as shown in Figure 1A. Hence, finding the optimal translation comprises finding occupied cells and local translations in those cells. This decomposition makes the encoding of the output invariant to translation and avoids manipulating large values to encode translations in the grid.

#### Architecture and learning procedure

We aim to solve the object detection problem stated above with a Machine Learning approach, trained on our dataset. Given a cryo EM density of size 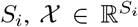 and its corresponding occupancy grid of size **S**, 𝒢 ^𝒳,S^, our network is a function 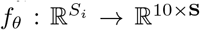 such that *ŷ* = *f*_*θ*_(𝒳) ∈ ℝ^10×S^ is our prediction for 𝒳. The prediction at a position *s* is denoted as *ŷ*^*k*^(*s*) ∈ ℝ^10^, where k = 1..10. The first dimension of this output, *ŷ*^0^ (·), is a prediction of the occupancy grid 𝒢^𝒳,S^. The nine other dimensions are predictions in each cell relative to the putative Ab contained in it. The details about both the exact role of each dimension as well as our training procedure are described below.

The architecture of our model *f*_*θ*_ is a 3D UNet [13] with a depth of 4. To enhance robustness, the network is trained using data augmentation, with the eight possible rotations over a grid and random cropping inside the input grid up to three cells. We used Adam optimizer over 1000 epochs. Hyper-parameters and exact architecture were not extensively tuned to avoid artificially boosting performance. Finer details about the ones we used can be found in Supplementary A.2.

#### Prediction of the right cells using optimal transport

For convenience, in this section, we drop indices and denote the ground truth occupancy grid 𝒢^𝒳,S^ as 𝒢. The first slice of our output *ŷ* is a prediction of the occupancy grid and hence, let us denote it as 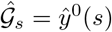. In order to make 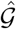 close to 𝒢, we will use two loss terms.

Because most grid cells are unoccupied, our prediction is very imbalanced. Hence, our first loss term is a weighted binary focal loss [38] that focuses on cells with *wrong* predictions :

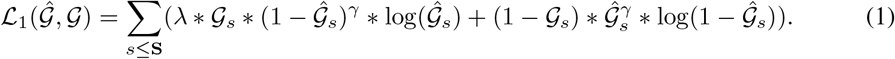

We used *λ* = 30 and *γ* = 4 in our experiments.

We observe that this focal loss ℒ_1_ does not consider the *distance* between our prediction and the ground truth. Predicting the neighbor pixel results in the same loss value as predicting the opposite side of the map. To address this issue, we add an *optimal transport* term based on Sinkhorn divergence to our loss [45]. After normalization, we can view 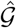 and 𝒢 as measures defined over the regular grid **g**. The optimal transport cost between those measures is defined as :

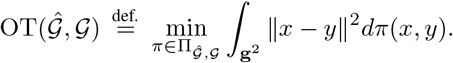

Where 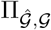 is the set of coupling measures on **g** with marginal 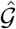 and 𝒢. The actual computation of this term is prohibitively expensive, hence we add a regularization term with weight *ε* > 0 and introduce :

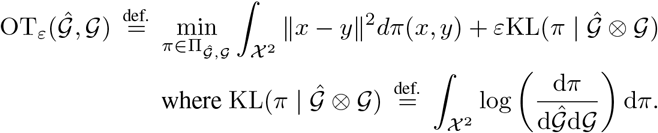

We notice that OT_0_ = OT and for any *ε* > 0, adding this εKL regularization makes the computation tractable. However, this regularization term also introduces the so-called *entropic bias* since for a measure *x*, OT_*ε*_ (*x, x*) > 0. Sinkhorn divergences were introduced to correct for this bias while remaining tractable, and that is what we use as a loss :

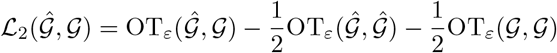

This term gives meaningful supervision to all voxels of our grids and depends on the distance to the closest occupied voxels. We used the *GeomLoss* package to compute it and refer to its corresponding paper [23] for a detailed discussion of the computation. The relevance of this loss is assessed in the results section.

#### Prediction within each cell

Let **A** be an antibody in our system and *s*_A_ its position in the grid. Beyond predicting the right grid cell, we also want finer grained prediction about its precise position in the cell 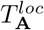, its orientation 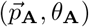 and its classification as a Fab or a VHH. Let 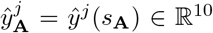 be the prediction at this position, we will introduce additional loss terms to capture these finer grained predictions. We emphasize that these will only be applied on grid cells containing an antibody. We use the notation 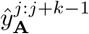 to denote the k dimensional vector obtained from the concatenation of 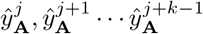.

To predict the right pose of **A** in the cell, we use eight values. The first three are used to predict the offset from the corner of the grid cell, 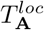. We use a mean squared error loss to learn these values, ℒ_3_. The following three values are used to predict 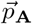 by directly predicting its coordinates. The corresponding loss, ℒ_4_, is composed of a dot product term to control the direction of the prediction along with a term to make this vector unit norm. Favoring unit norms avoids numerically unstable normalizations. Finally, the remaining two values are used to predict the angle *θ*_A_. Instead of directly predicting the angle, we aim to predict *u*_A_ ∈ ℝ^2^, the vector of polar coordinates (1, *θ*_A_). This formulation avoids singularities and was shown to be beneficial when predicting angles [31]. Hence, ℒ_5_ has a similar form than ℒ_4_ in two dimensions to predict *u*_A_. We end up with the following losses:

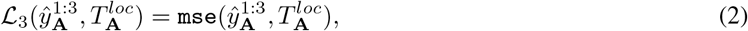

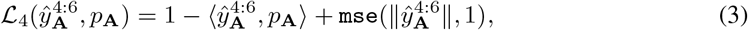

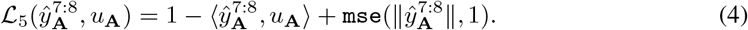

Finally, as we use a single model for both Fabs and VHHs, we have a term that represents the probability that the object contained in the grid cell is a VHH and not a Fab. Let *δ*_*n*_(*x*) be the indicator function for VHHs (one if VHH else zero). We again construct a weighted focal loss, with a weight of *λ*_*n*_ = 1000/400 corresponding to the ratio of VHHs to Fabs, yielding the last loss,

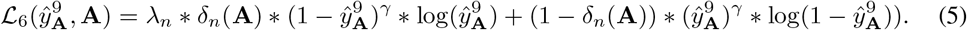

To train our network we use a weighted sum (*λ*_*s*_ = 0.2) of previous loss terms as the final loss. We sum the first two global ones and the sum of the four others over each antibody in our system:

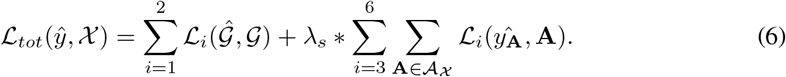

#### Post processing

A well-known problem with object detection is the possibility that the network predicts overlapping objects. Supposing occupancy grid cells edges were 3Å long, adjacent cells could not both contain a Fab. Hence, high values for adjacent cells typically amount to the detection of the same underlying object. Non Maximal Suppression (NMS) algorithms are used to discard such redundant predictions. Starting with our grid *ŷ*_0_, we want to obtain a list of the distinct local minima. In this paper, we used an approach based on `persistence diagrams` [9], implemented with *cripser*[32, 57]. Simply put, we decrease a threshold probability value from the maximum value of our grid *ŷ*_0_ and keep track of cells above this threshold. When the value of a cell goes over the threshold, either it has no neighbors belonging to a visited connected component, giving *birth* to a new one, or all neighboring components are merged into the one with lowest initial values and merged ones *die*. The difference between the values of death and birth are called lifetimes. We return connected components sorted by lifetimes.

This procedure takes into account both the *value* of a minimum and its location with respect to other minima. If we suppose that the number of objects to find is known, we keep proceeding until this number is reached (and refer to this setting as and `num`). Otherwise we use a lifetime threshold and retain all predictions above this threshold. We use a threshold of 0.2 that was found to work best on the validation set (see Supplementary A.4). We will denote this setting `thresh`. Interestingly, this procedure allows us to automatically detect the numbers of Abs in a density, without any prior information.

From our density map, we now have a list of predicted values. For each of those, we choose a template based on the classification in Fabs and VHHs and move this template to the predicted location and pose. We save the result as a file in PDB format.

## Acknowledgments and Disclosure of Funding

This research was supported by Sanofi. V. M. and M. O. are supported by DataIA, ANR AI Chair AIGRETTE and the ERC Starting Grant EXPROTEA. C.R. and H.M. are employees of Sanofi and may hold shares in the company.

## Appendix

### A Methods

#### A.1 Proof of the validity of the decomposition

Following our notations, 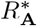 is a rotation that aligns **A** onto **T**. We want to show that 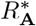 can be parametrized by the transformation of *p*_T_ into *p*_A_ along with a rotation around *p*_T_.

By construction :

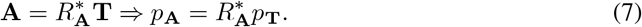

Let *M* be a rotation matrix of axis *p*_A_ × *p*_T_ and of angle *ρ* the angle between these vectors, so that it transforms one into the other. We can write :

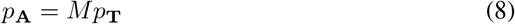

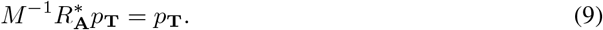

Therefore, 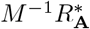 is a rotation around the axis *p*_T_ by some angle *θ*, and we can write :

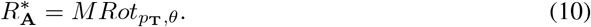

#### A.2 Detailed architecture

We used a 3D Unet architecture. First let us define a `conv_block`(i,o) as three series of convolutions (with kernel size three and stride one), batch normalization and PreLu activations. The first convolution goes from i to o channels and the two others have o as a fixed number of channels.

In the encoding stage, we use a series of four `conv_blocks` of successive dimensions (1, 32, 64, 128, 256), alternated by a max pooling of kernel size 2 and stride 2.

In the decoding stage, we apply a transpose convolution (256, 256) with kernel size of 3 and stride 2 to the output of the encoder. We concatenate the result with the output from the third encoding `conv_block` to get 384 channels. We then run a `conv_block`(384, 128) on this concatenation.

Finally, we apply a last `conv_block`(128, 128) with only two series on the resulting tensor and replace the third one by a linear layer going from dimension 128 to dimension 10. The classification output dimensions are followed by sigmoid activation.

#### A.3 Modifications to the SabDab database

We removed a few non canonical systems, such as fusion proteins or `8gq5` that contains 16 VHHs arranged in an unusual way and the obsolete `7ny5`. We then fixed edgy situations, often encountered when VHHs and Fabs are present in the system.

- When querying the Fabs, some chains were wrongly considered separate : `7YVN chain HI 7YVP chains IJ, 8HHX chains HI, 8HHY chains FW-GI, 8DXS chains FI-GJ, 7XOD chains RS-UV-XY, 7MLV chains KF, 6XJA (missing chain AB and missing antigen)`
- VHHs were present when querying Fabs. They were annotated as having just one chain and no antigen partner : 7zlg-h-i-j chain K bound to L, 8hii-j-k chain N bound to L, 7xw6 chain N bound to AB, 7sk7 chain K bound to C, 7sk5 chain E bound to C, 7zyi chain K bound to L, 7xod chain T-W-Z bound to S-V-Y, 6ni2 chain A bound to B-V, 7wpe chain W-Z bound to V-Y, 6ww2 chain K bound to L, 7jhh chain N bound to L, 7ul3 chain C bound to A, 7tuy chain K bound to L
- Fabs were present when querying VHHs (often interacting with this VHH) : `7pij chain HL bound to N, 7wpd chain XY bound to A, 7wpf chain RS, UV, XY, 7wpf chain AC-HL bound to EN, 7php chain HL bound to N, 7m74 chain HL bound to A`
- VHHs with no antigens were present in the VHH file (actually always bound to a Fab) : 7pij K bound to L, 7wpd Z bound to Y, 7nkr F bound to A, 7wpf T-W-Z bound to S-V-Y, 7wpf chain K-F bound to L-C, 7php-7m74-7jhg chain K, N, N bound to L, 7nis-7nk1-7nk2-7nk6-7nkc-7nj4-7nka chain F bound to A/B-A-A/B-A/B-B/C-A-A

#### A.4 Finding the optimal threshold

We present the error in the number of Abs detected for different thresholds in Figure S.1. Throughout datasets, persistence diagrams seem to give a better estimate. The overall best value is achieved for a threshold of approximately 0.2 and is not very sensitive to this threshold.

**Figure S.1:**
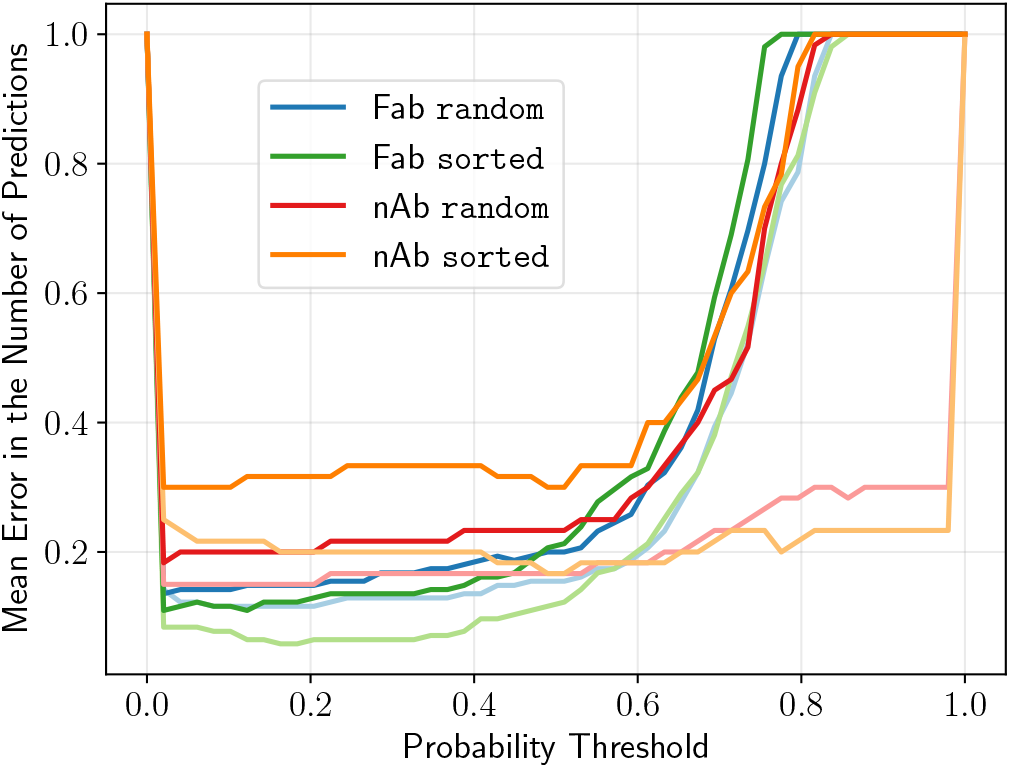
Finding the best threshold for selecting the right number of system. x-axis represents the threshold value. y-axis represents the mean absolute error in the number of predicted systems over the data set. Different colors represent different data sets. Darker lines represent a naïve approach for NMS while the lighter ones are computed using persistence diagrams.

### B Results

#### B.1 Fab detection performance

We compute the distances from prediction to ground truth obtained from our tool and `dock_in_map`. For a nicer visualization, we cap all distances to 20Å. We also count failed systems and errors as 20Å predictions so that the bar at 20Å represents all failures of a tool. We show the result in Figure S.2

**Figure S.2:**
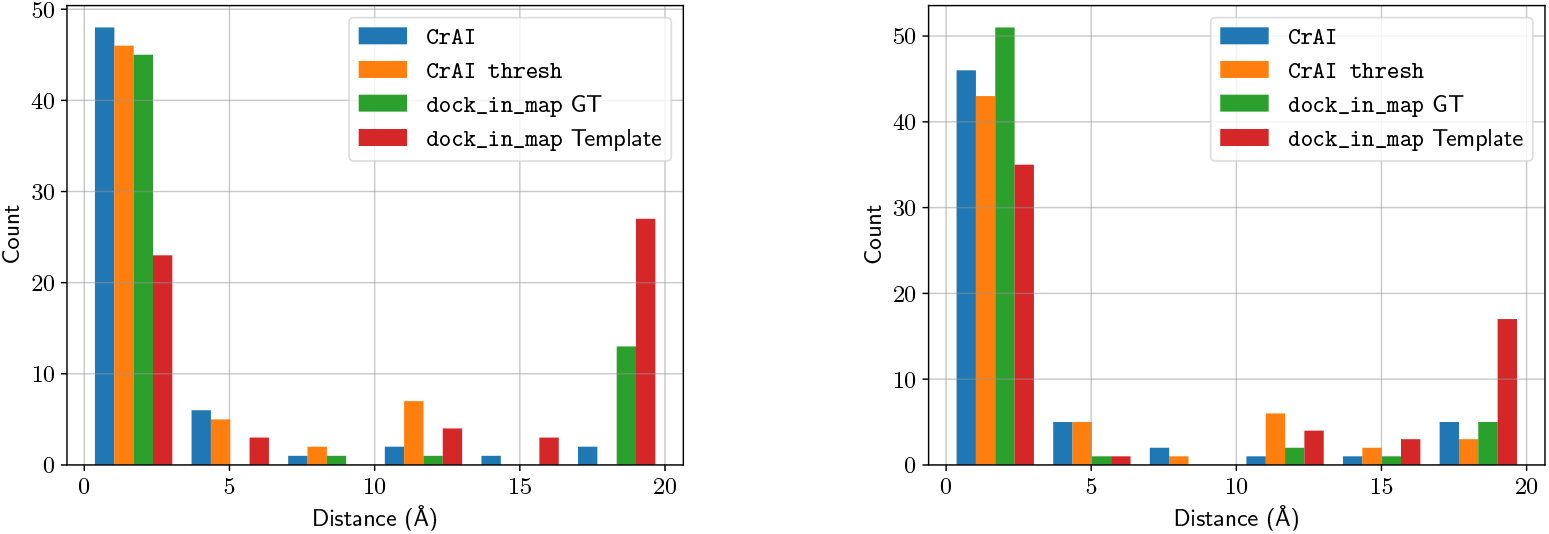
Capped histogram of distances between predictions and experimental structures on Fabs. `random` is on the left and `sorted` on the right.

As can be seen, our tool widely outperforms `dock_in_map`. Under the hit rate metric, we have the following performances (by order of tools) : `random` : 94.1, 88.8, 62.2, 19.4 `sorted` : 97.3, 93.9, 71.1, 8.5.

#### B.2 Dependence in resolution

We concatenate the predictions for all our systems and scatter them as a function of the resolution of the maps. We cap the predicted distances to 10Å and add a best fit curve. We show the result in Figure S.3. We see a moderate dependency in resolution, and many successful ones for resolutions above 4. In contrast, `dock_in_map` exhibits a strong dependency to resolution.

**Figure S.3:**
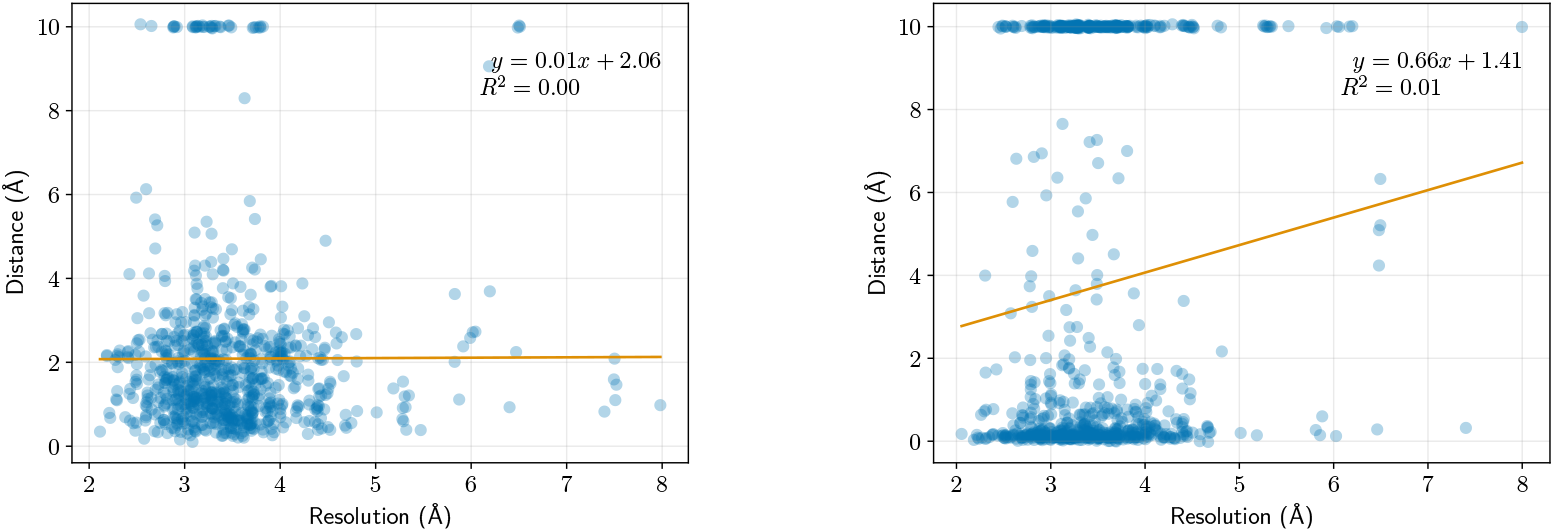
Scatter plot of the performance of `CrAI` (left) and `dock_in_map` (right) as a function of the resolution. x-axis is the resolution of the maps. y-axis is the distance between a prediction and its corresponding system, capped to 10Å for failed systems.

#### B.3 Failed examples

**Figure S.3:**
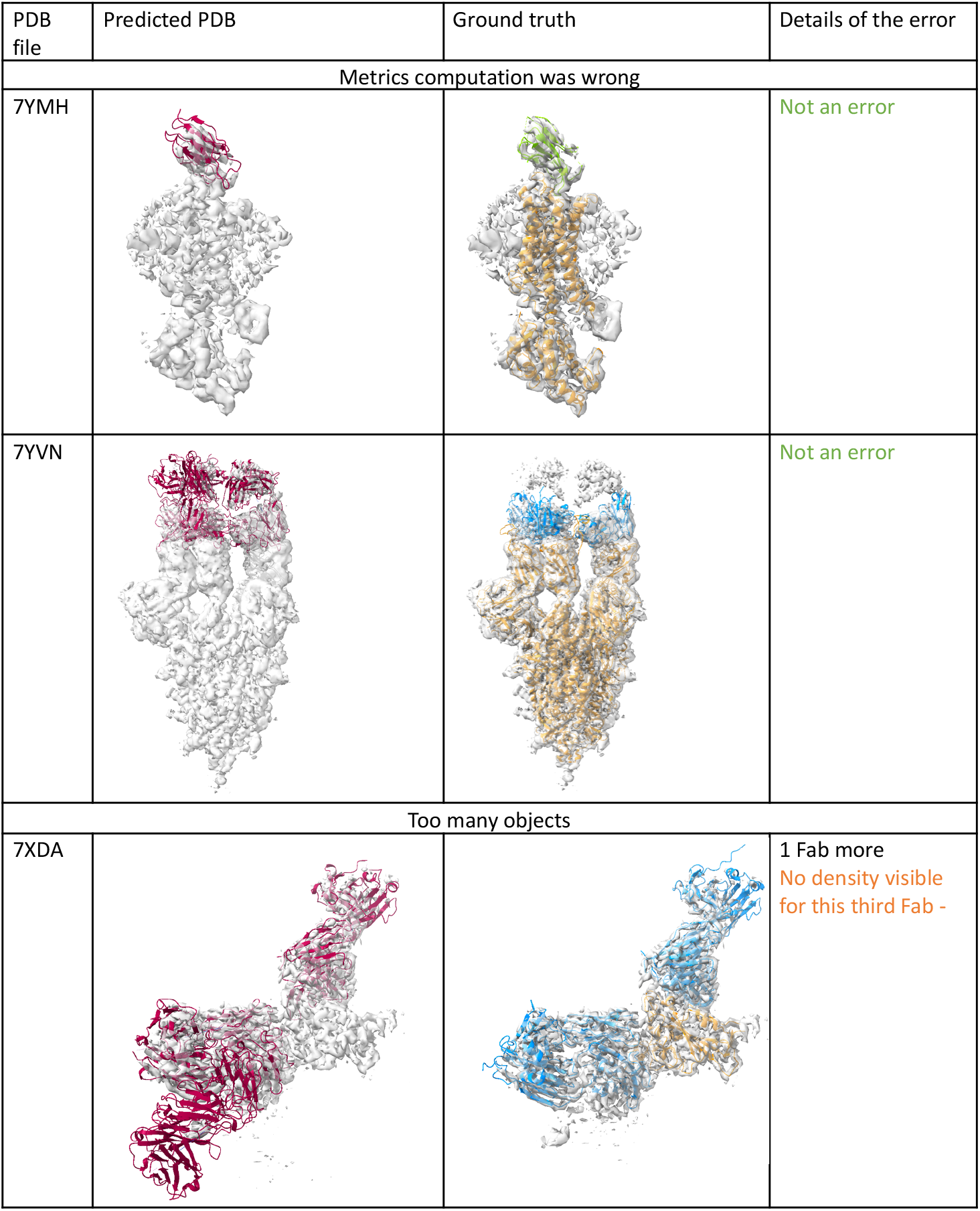
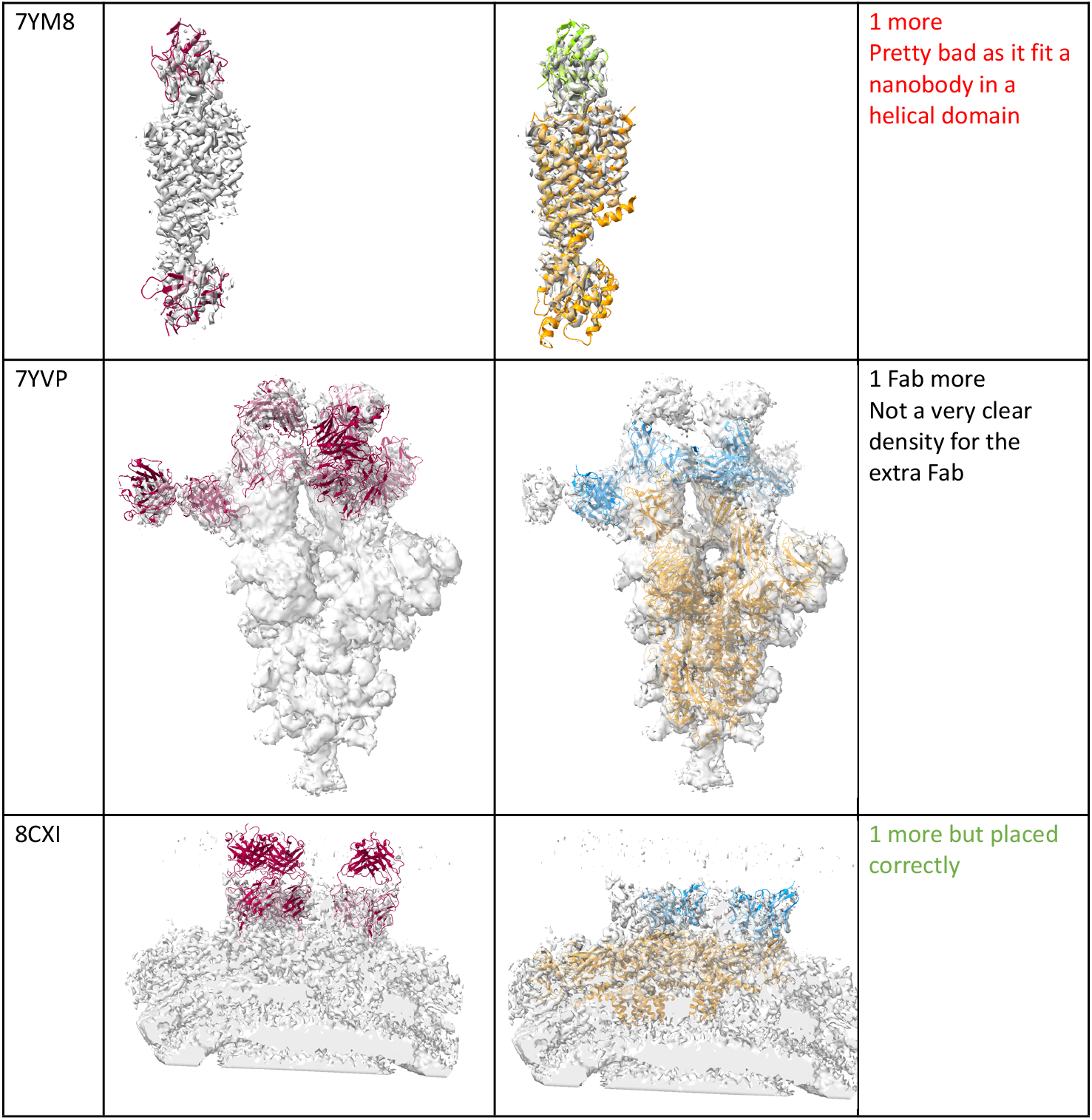
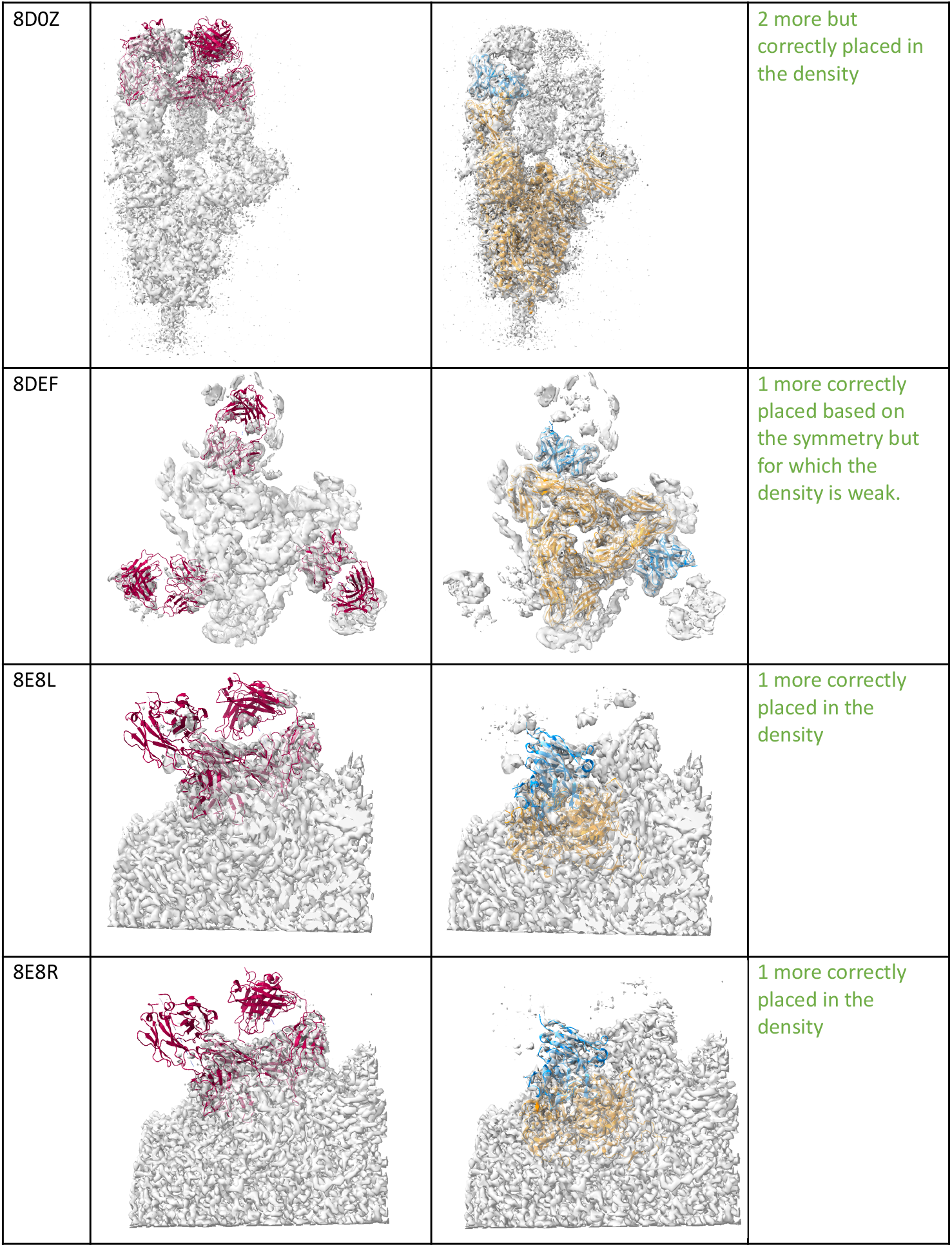
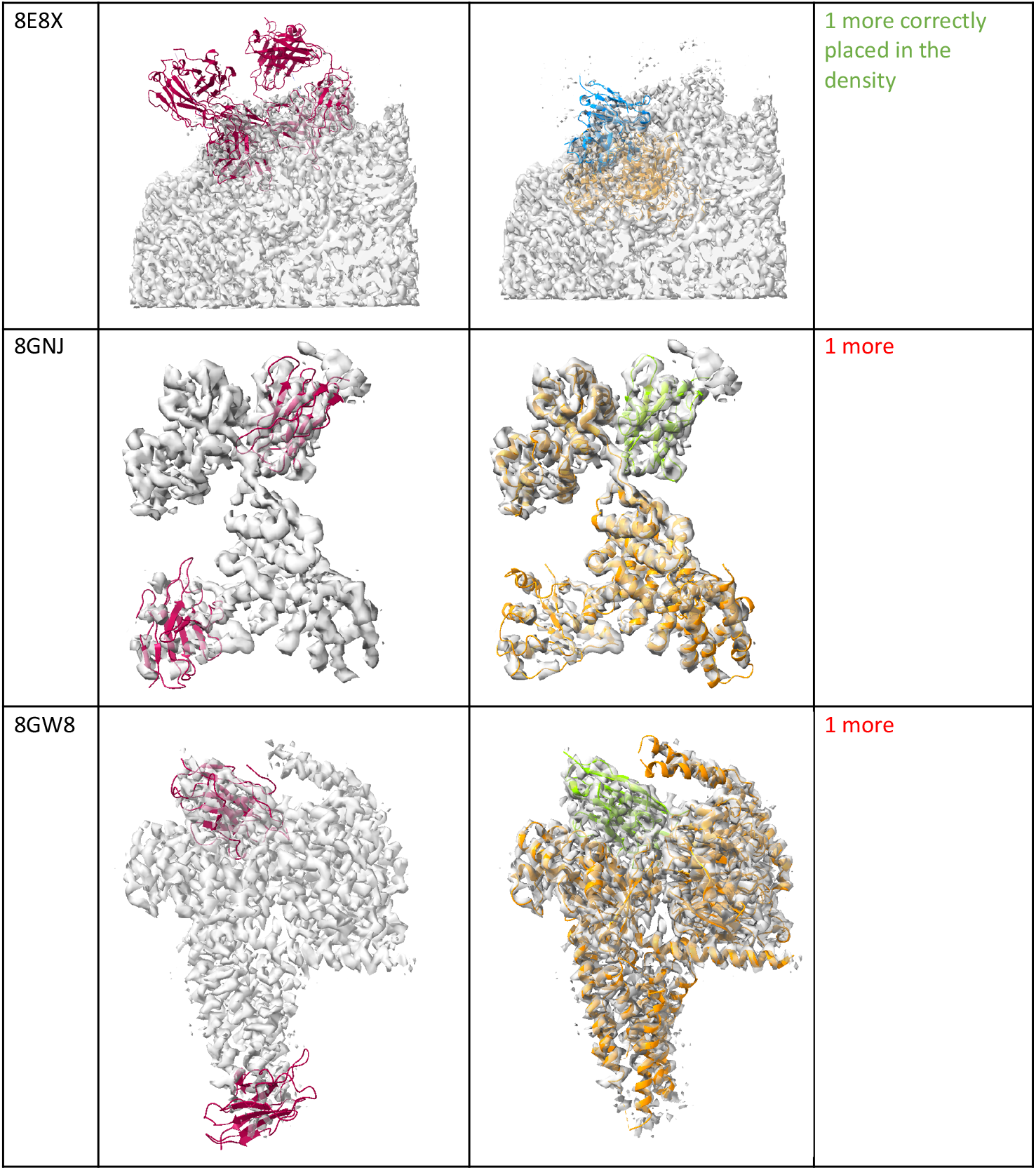
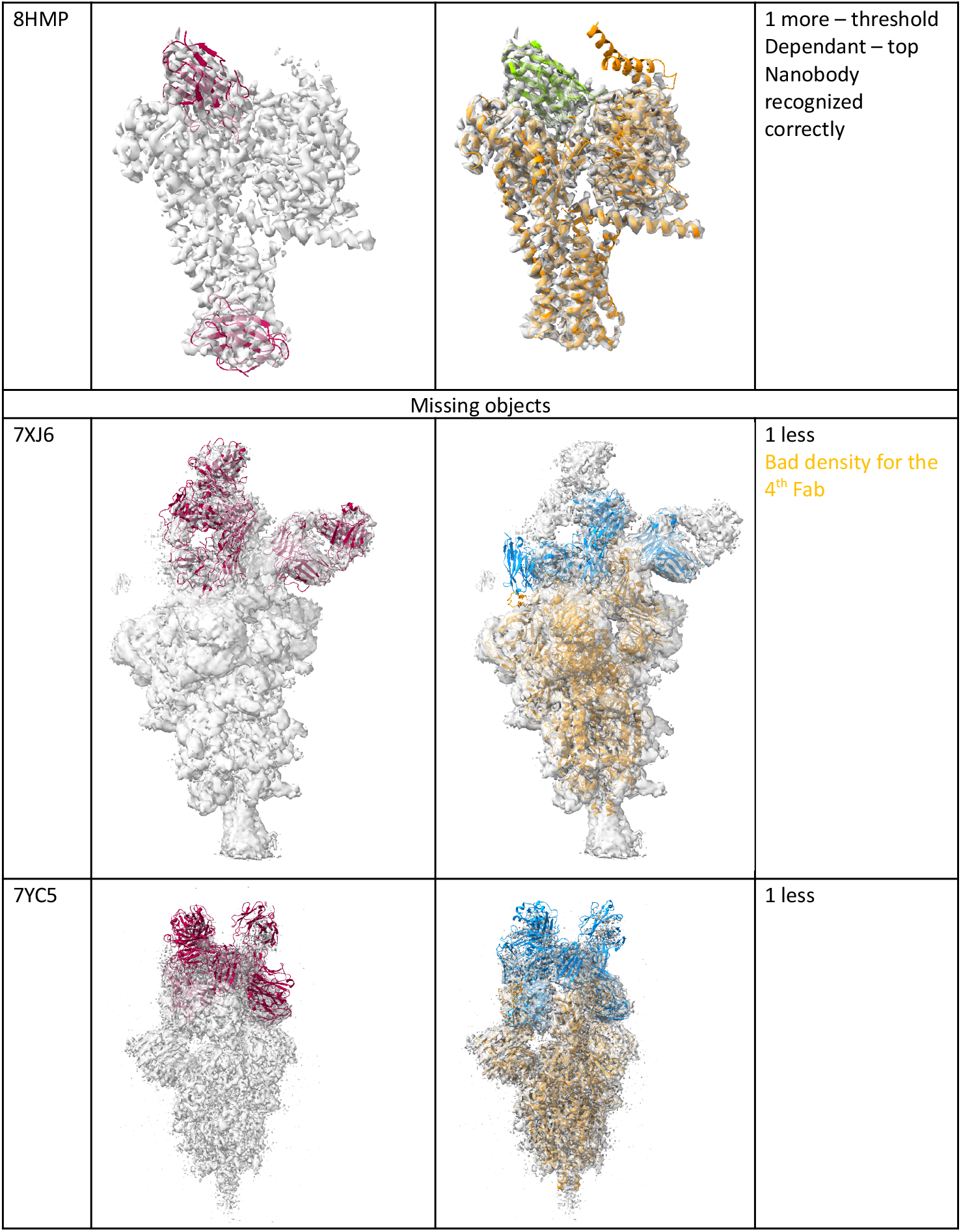
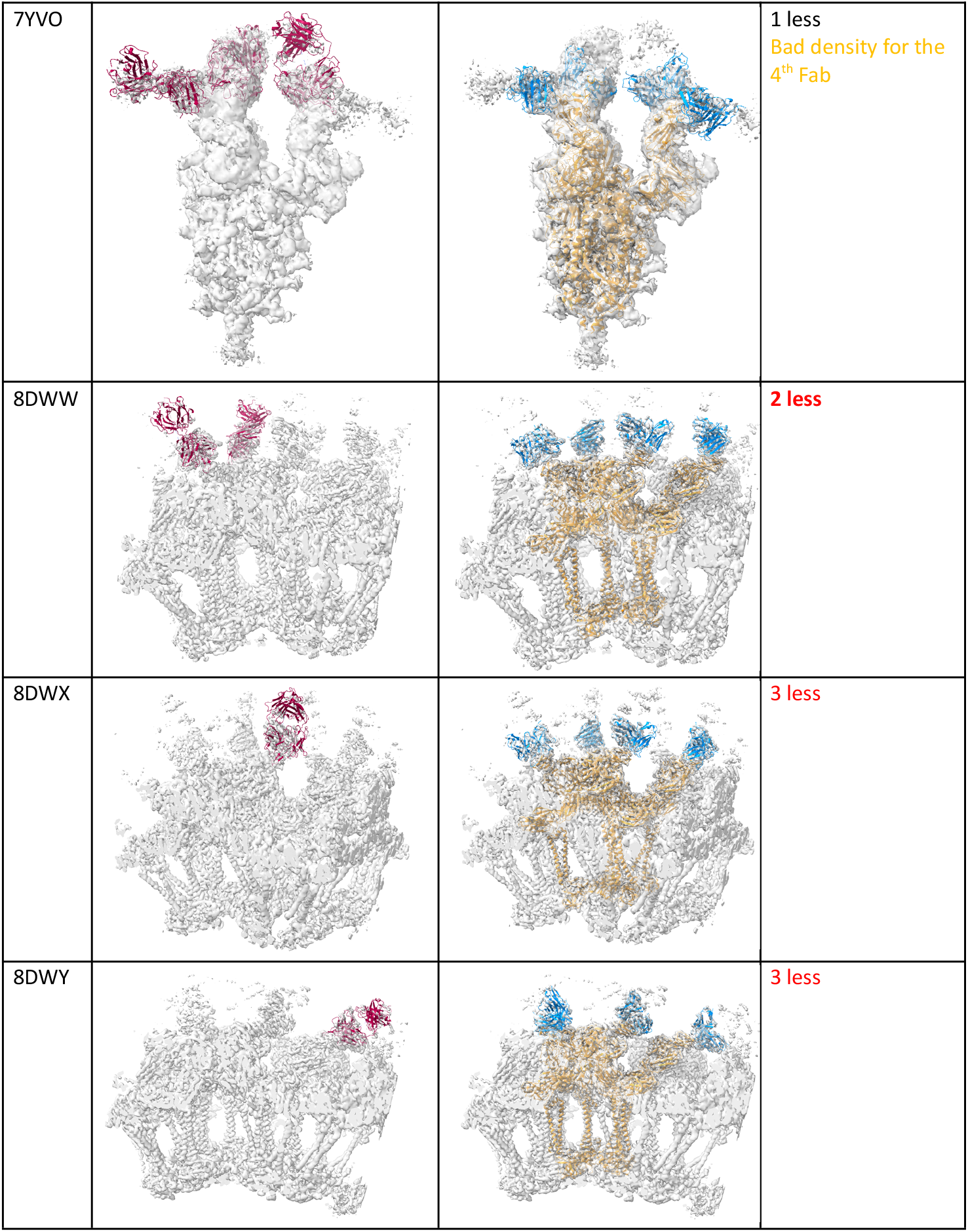
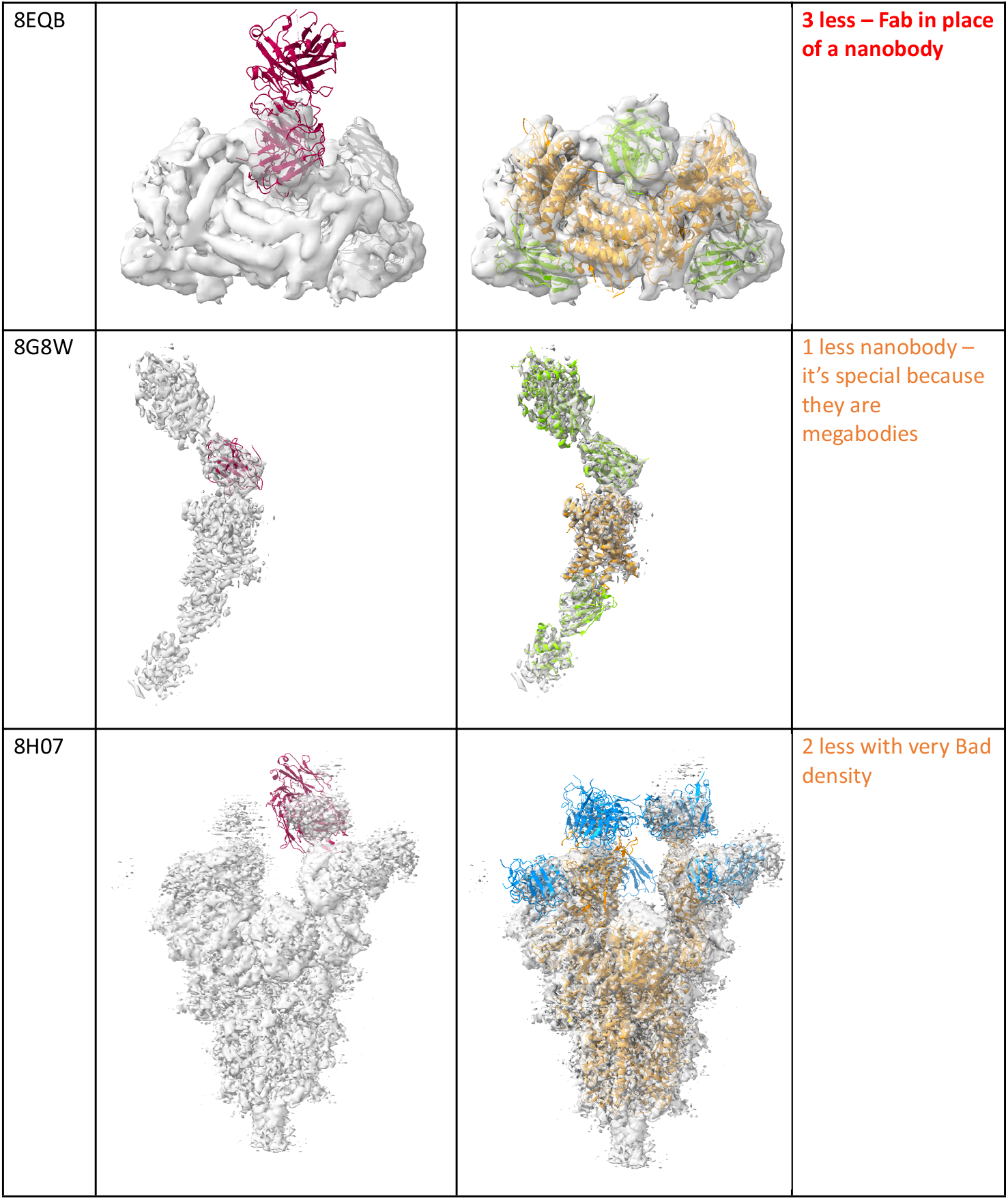
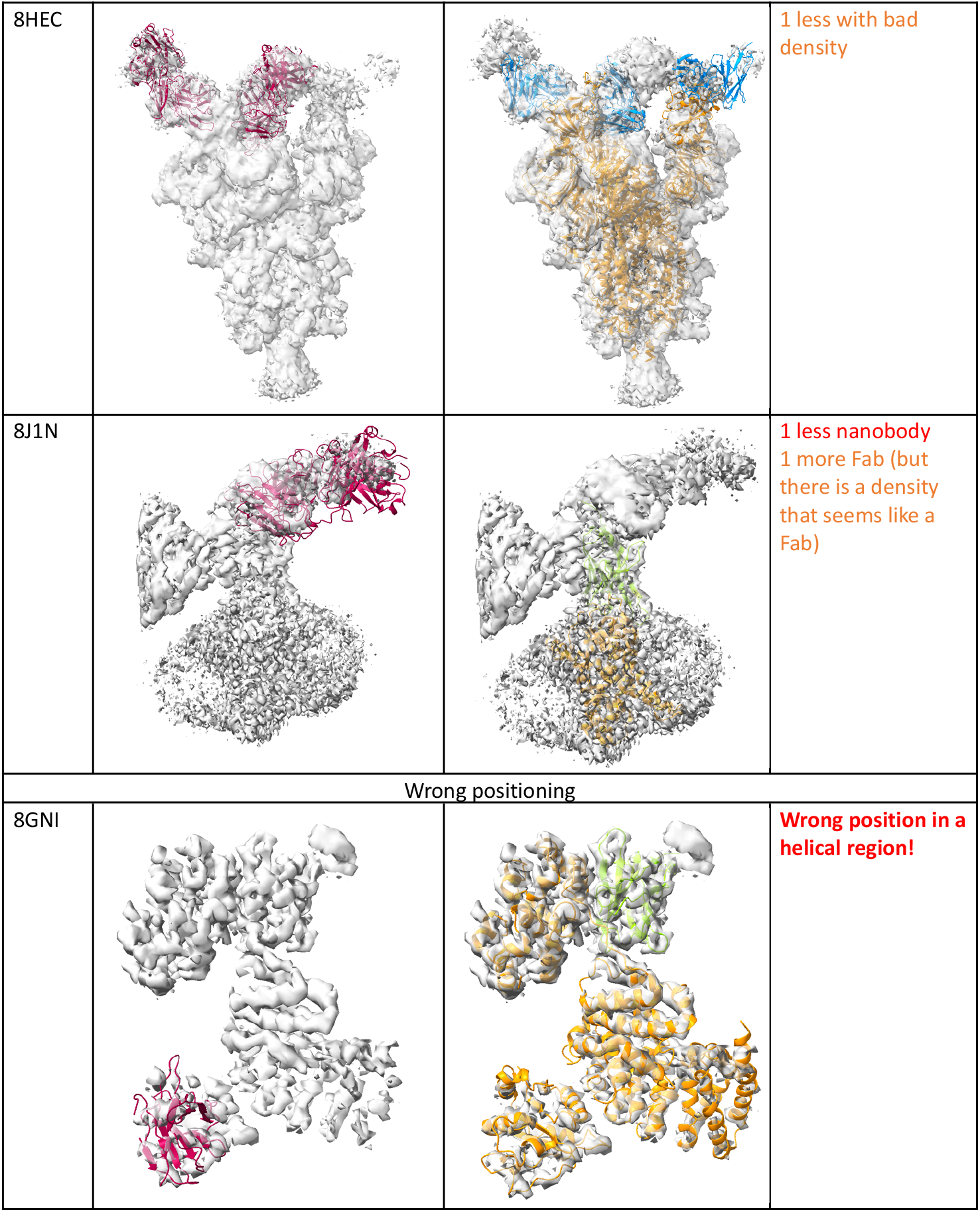
All failures, split by error type

## Footnotes

1 github.com/Sanofi-Public/crai and cxtoolshed.rbvi.ucsf.edu/apps/chimeraxcrai

